# The Landscape of Enteric Pathogen Exposure of Young Children in Public Domains of Low-Income, Urban Kenya: The Influence of Exposure Pathway and Spatial Range of Play on Multi-Pathogen Exposure Risks

**DOI:** 10.1101/388140

**Authors:** Danielle Medgyesi, Daniel Sewell, Reid Senesac, Oliver Cumming, Jane Mumma, Kelly K. Baker

**Affiliations:** Department of Occupational and Environmental Health, University of Iowa, Iowa City, Iowa, United States; Department of Biostatistics, University of Iowa, Iowa City, Iowa, United States; Department of Disease Control, London School of Hygiene and Tropical Medicine, London, United Kingdom; Department of Community Nutrition, Great Lakes University of Kisumu, Kisumu, Kenya

## Abstract

**Background:** Young children are infected by a diverse variety of enteric pathogens in low-income, high-burden countries. Little is known about which conditions pose the greatest risk for enteric pathogen exposure and infection. Young children frequently play in residential public areas around their household, including areas contaminated by human and animal feces, suggesting these exposures are particularly hazardous.

**Objectives:** The objective of this study was to examine how the dose of six types of common enteric pathogens, and the probability of exposure to one or multiple enteric pathogens for young children playing at public play areas in Kisumu, Kenya is influenced by the type and frequency of child play behaviors that result in ingestion of soil or surface water, as well as by spatial variability in the number of public areas children are exposed to in their neighborhood.

**Methods:** A Bayesian framework was employed to obtain the posterior distribution of pathogen doses for a certain number of contacts. First, a multivariate random effects tobit model was used to obtain the posterior distribution of pathogen concentrations, and their interdependencies, in soil and surface water, based upon empirical data of enteric pathogen contamination in three neighborhoods of Kisumu. Then, exposure doses were estimated using behavioral contact parameters from previous studies, and contrasted under different exposure conditions.

**Results:** Multi-pathogen exposure of children at public play areas was common. Pathogen doses and the probability of multi-pathogen ingestion increased with: higher frequency of environmental contact, especially for surface water; larger volume of soil or water ingested; and with play at multiple sites in the neighborhood versus single site play.

**Discussion:** Child contact with surface water and soil at public play areas in their neighborhood is an important cause of exposure to enteric pathogens in Kisumu, and behavioral, environmental, and spatial conditions are determinants of exposure.

## Introduction

Children living in low income countries with poor sanitary conditions experience an average of 4 to 8 diarrheal episodes per year between birth and 2 years of age,(1) demonstrating that they are chronically exposed to enteric pathogens beginning in the first year of life. Furthermore, recent studies have highlighted wide diversity in the microbial etiology of early childhood (<5 years of age) enteric infection in such settings, suggesting that they are exposed to a variety of pathogenic organisms in the first years of life.(2–9) Little is known about the rate with which children are exposed to and acquire enteric infections over time. While diarrheal incidence may suggest an exposure rate of up to 4 to 8 pathogen types per year, many infections are asymptomatic and go undetected without extensive diagnostic profiling,(10) meaning that diarrhea symptoms are likely an underestimate of how often children acquire new infections. Diarrhea rates may even further underestimate how often children are exposed to pathogens, but remain uninfected due to insufficient exposure dose, lack of pathogen viability, host acquired immunity, or other mediating conditions. Adding to this complexity, co-infection of individual children by two or more types of pathogens – regardless of symptomology – is common.(2, 3) Co-infection, even in the absence of diarrhea, is associated with greater risk of environmental enteric dysfunction (EED), undernutrition, and re-infection by a new pathogen, perpetuating the cycle of disease.(4) Understanding which exposure pathways contribute most to multi-pathogen exposure of children could improve the prioritization of interventions that reduce early childhood enteric disease incidence.

Wagner and Lanoix’s “F-diagram” conceptualized the routes of fecal-oral disease transmission according to the properties of environmental materials (drinking water, food, soil, etc.) that can be contaminated by feces and ingested by humans.(11) However, there is limited research on how exposure varies across exposure pathways, particularly with respect to the rates at which children experience multi-pathogen exposure and infection. Existing comparisons of exposure pathways have relied on fecal indicator bacteria concentrations,(12–17) or pathogen-specific risks(18–20), both of which have major methodological limitations in describing the overall risk of enteric pathogen exposure across pathways. Different types of pathogens have been frequently detected in households of India and Tanzania and public play areas in Kenya, revealing that exposure to pathogens in private and public settings is likely.(21–23) Our group has further shown that soil and surface water from public areas where children play in Kisumu are often contaminated *simultaneously* by multiple types of pathogens,(23) revealing that children ingesting soil or water at some public sites could ingest multiple types pathogens. Only one report to our knowledge has examined exposure from the perspective of ingesting multiple types of pathogens, rather than presence/absence of an indicator.(24) But, the modeling approach summed the individual probabilities of exposure to each type of pathogen from South African surface waters, rather than accounting for interrelatedness of pathogen contamination across sampling locations or exposure pathways. Since multi-pathogen contamination varies across location, exposure models must account for possible pathway- or location-specific differences in multi-pathogen contamination and transference.

The overall risk of exposure to enteric pathogens may be fundamentally different across exposure pathways and across location of exposure. Some differences may be caused by physical properties of the environmental fomites. For example, eating soil from the ground may be more hazardous than ingestion of household drinking water because humans or animals may defecate directly on the ground whereas water is more likely to be protected and treated for safety.(12, 25, 26) Some exposure differences may be related to child behavior. Young children typically have high rates of contact with soil and objects,(27–31) and occasionally surface water,(32, 33) and frequently place their hands in their mouth with no handwashing in between.(33, 34) This results in frequent indirect ingestion of trace amounts of soil, and perhaps surface water. Geophagia (direct ingestion of handfuls of soil) among young children(25, 26, 33, 35) and drinking from surface water also occur, (33) albeit less frequently than hand-to-mouth behaviors. Yet, the greater volume of material ingested by these infrequent behaviors could result in greater pathogen doses than what is ingested via hand-to-mouth behaviors. Finally, spatial variability of young children’s play in neighborhoods could influence the dose and diversity of pathogen exposure. Many children play outside the household unattended, while others have self-or guardian-driven limitations that restrict distance away from the household and acceptable areas for play.(33) In settings where the landscape is often contaminated by feces from many humans and animals, the children who play in a constrained spatial area (i.e. near their household) may experience lower risks for feces exposure than children who roam across a larger spatial area and play at a variety of locations throughout the course of a day, especially if those locations are public areas used for feces disposal. More knowledge on how child behavior, type of environmental fomite, and spatial range of child play influences enteric pathogen exposure risks is needed for prioritizing interventions.

This study proposes an innovative exposure assessment approach that utilizes information on spatial variability in enteric pathogen detection and co-detection across public play areas in a typical low-income, fecally-contaminated setting to explore the relative importance of different environmental, behavioral, and spatial conditions in pathogen exposure of young children. The first objective of this study was to measure how increased frequency of child contact with soil or surface water and the volume ingested (indirect vs. direct ingestion) by children in public play areas influences the ingestion dose of enteric pathogens, and the probability of exposure to one or more enteric pathogens. Second, we compare how constrained (child plays at one public residential location) versus neighborhood (free roaming) spatial range of play for children influences pathogen dose and probability of multi-pathogen exposure. This modeling approach could be adapted to include a variety of setting-specific information on child behaviors and environmental conditions to better assess the relative contribution of various exposure pathways to child infection.

## Materials and methods

### Study design

Observational assessment and environmental microbiology data on public sites in Kisumu, Kenya has been described previously.(23) In brief, 166 total public sites in three peri-urban neighborhoods of Kisumu were randomly selected for a cross-sectional observation and environmental sampling study. A site was defined as all public area (private households and businesses excluded) falling within a 25-meter radius. Enumerators documented environmental conditions of the site such as landscape features, condition of public or communal latrines, and indicators of human open defecation or unsafe disposal of excreta. Presence and type of domestic animals and their feces were recorded. During rapid observation (~10-15 minutes per site), enumerators recorded whether children approximately <5 yrs were observed at the public site and their behaviors that would result in hand or mouth contact with environmental fomites (touching soil, surface water, animals, or objects on the ground, swimming, eating food, eating dirt, mouthing hands). Approximately five grams of soil was collected at every site, and 10 mls of surface water was collected if present. Samples were analyzed to quantify the concentration of enterococcus indicator bacteria and 19 types of common enteric pathogens using qRT-PCR. Four resamples were collected at seven randomized sites in each neighborhood to account for anticipated variance in pathogen distributions at public sites.

During rapid observation at least one child was observed to play at 54% of public residential sites and engaged in behaviors such as hand contact with soil and surface water, hand-to-mouth contact, and geophagy, validating that observed behaviors in Kisumu neighborhoods are consistent with extant literature (25–27, 32, 33, 35, 36) and are relevant pathways for pathogen exposure. Of all public sites where children <5 yrs were observed in Kisumu, 93% of these sites were residential areas with mostly permeable or unpaved surfaces, versus non-residential sites. This study uses microbial data from 125 soil samples and 34 surface water samples from 125 residential public sites - the locations where children are most likely to spend time. Of these, 30 soil samples and 3 surface water samples were randomly selected re-samples.(23)

To ensure sufficient knowledge about the distribution of pathogen concentrations and to obtain numerical stability in the statistical estimation algorithms, pathogens detected in fewer than 5% of the soil and surface water samples were excluded from the model. Of the 19 pathogens tested during environmental sampling, concentration data for 6 pathogens (Table S1) were eligible for inclusion in the model: *Cryptosporidium spp., Giardia*, human adenovirus 40/41, Enteropathogenic *E. coli* (EPEC *bfpA* and/or *eae*), Enterotoxigenic *E. coli* (ETEC *est* and/or *elt),* and Enteroaggregative *E. coli* (EAEC *aiicA* and/or *aatA).(23)* If there was a positive detect for more than one bacterial gene marker, and concentrations (C_p_) varied between gene markers, concentrations of ETEC-estA, EPEC-*bfpA*, and EAEC-*aatA* were prioritized over concentrations of *ETEC-eltB,* EPEC-eaeA, and EAEC-aaiC, respectively, based on etiological importance in pediatric diarrheal disease.(2, 3) Although the dataset was restricted to 6 of 19 pathogens measured, the number of soil and surface water samples with at least one positive detect (138/159 samples) did not change, indicating that these 6 pathogens collectively are sensitive indicators for the presence of pathogen contamination in the environment in this setting.

### Statistical analyses

The statistical analyses aimed to estimate the dose distributions of each pathogen type by environmental fomites type (soil vs. surface water), by contact type (indirect hand-to-mouth vs. direct geophagy or drinking), frequency of contact, and spatial range of exposure (site-restricted vs. random selection across the neighborhood). All analyses were conducted using RStudio version 3.4.2 (Boston, MA, USA). A Bayesian framework was employed to obtain the posterior distribution of pathogen doses for a certain number of contacts, denoted as *D* (*k*) for contact events, i.e., the distribution of *D*(*k*) implied by information provided by both our data and previous studies.(14, 26, 31, 37–40) The posterior distribution yields point estimates and credible intervals for the parameters of the pathogen concentration distribution, denoted as *θ*, for soil and water samples.

There are two parts to the modeling framework. The first part uses environmental microbiology data (Table S1) to estimate the distribution of each pathogen concentration in soil and surface water.(23) The second part combines the concentration distribution of part one with contact fate parameters provided from previous studies (Table 1) to estimate the exposure pathway-specific dose distribution by fomites type, contact type, and behavior frequency. The posterior distribution of interest, namely that of *D*(*k*) and *θ* given the observed data and information from previous studies, can be decomposed to clearly reveal these two components of the statistical model:

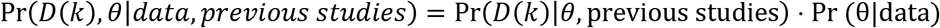

**Table 1.**
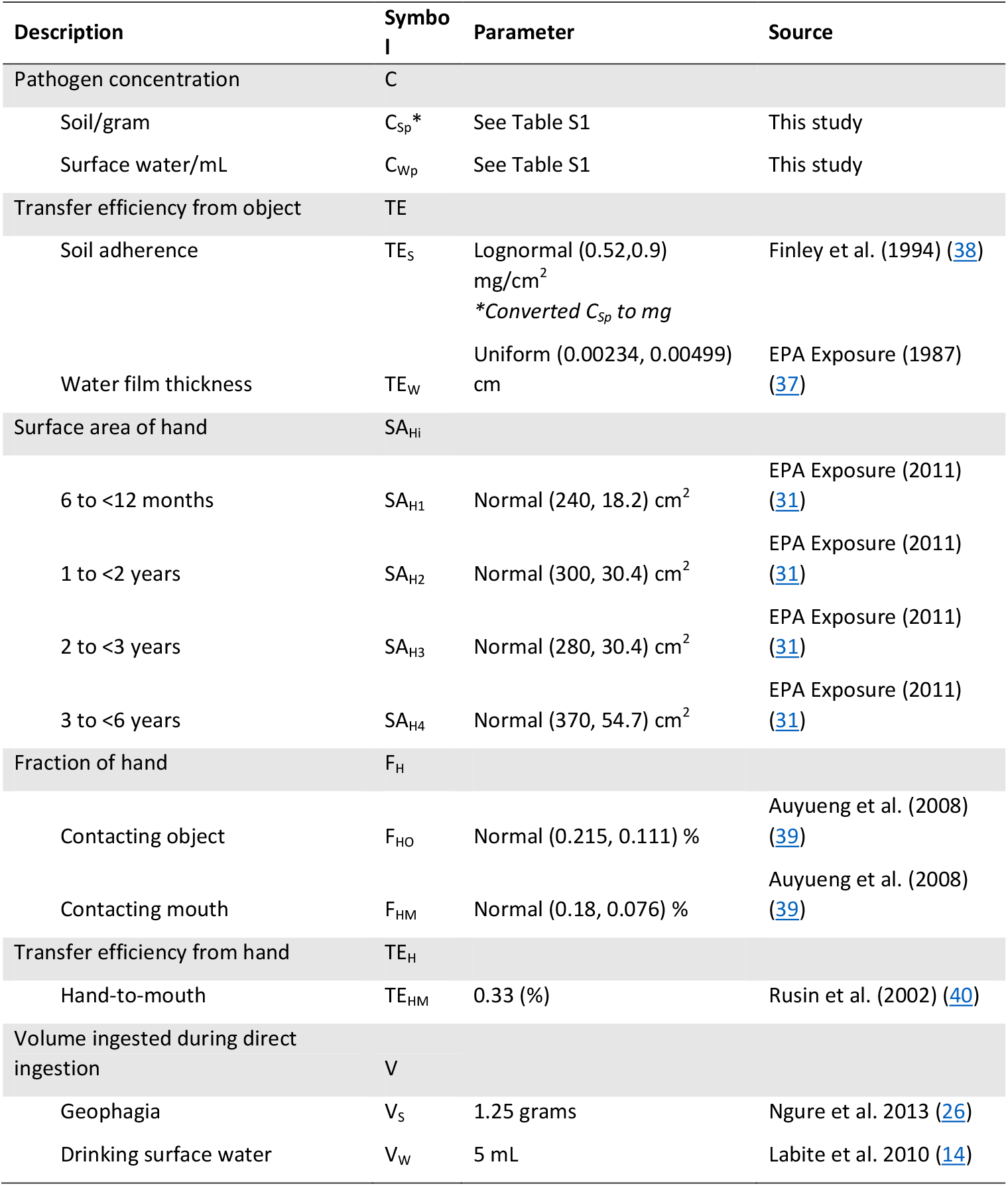
Description of variables and parameters used to estimate distribution of indirect and direct doses.

In implementation, we estimated this posterior distribution via a Monte Carlo approach on the joint posterior distribution augmented with the pathogen concentration corresponding to the events. That is, we may first obtain a sample of *θ* from the marginal posterior distribution given the data then draw pathogen concentrations for the current value of *θ*, and finally, given those concentrations and information obtained from previous studies on exposure pathways, draw *D*(*k*).

### Distribution of pathogen concentrations in soil and surface water

Several challenges arose in estimating the parameters *θ* for the pathogen concentration distributions. First, there was left censoring caused by methodologically-constrained lower limits of detection (Table S1). Second, there were two important sources of dependency in the data—that which occurs due to the correlations between the different pathogens, and that which occurs due to resampling within the 25-meter radius area of as an individual site.

To handle data challenges, we fit a multivariate random effects tobit model to the log transformed concentration data. The first source of dependency in the data was accounted for by modeling all pathogens jointly rather than running many univariate analyses. It was also important that we not neglect to account for the latter type of dependency described above, as the spatial patterns of young children playing in neighborhoods could influence the dose and diversity of pathogen exposure in public areas. Thus, the proposed random effects included in our multivariate tobit model account for this spatial dependence. See S1 Appendix for details on this statistical model.

For each environmental sample type, samples of *θ* were obtained from the posterior distribution using a Gibbs sampler. From these samples, posterior draws of pathogen concentrations of new events were drawn from a multivariate normal distribution parameterized by the draws of *θ*. See the S1 Appendix for details on the Gibbs sampling algorithm.

### Exposure pathway-specific dose distribution

A theoretical model was developed to estimate and compare the dose and diversity of enteric pathogens ingested by young children via indirect and direct exposure to soil and surface water at public play areas. The contact frequency was held at a constant rate, ranging from a minimum of 1 to a maximum of 10 contacts, for pattern comparison purposes, so behaviors in this model are not weighted to account for the likelihood of engaging in the behavior and the rate of contact given a child plays in a public area for a specified time span.(33) Therefore, the results are not cumulative estimates of actual child exposure, but represent possible exposures given a range of possible conditions. To obtain a posterior sample of the final dose for each set of conditions, the concentrations drawn previously were multiplied by fate parameters, each drawn from a random distribution to account for the inherent variability in such occurrences, and then the doses ranging from 1 to 10 were summed. The spatial assumption determined whether or not the contacts were correlated. The formulas used to estimate the dose distribution from indirect and direct contact with soil (1) and surface water (2) are:

1. **Soil**

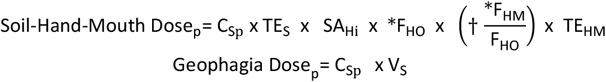
2. **Surface Water**

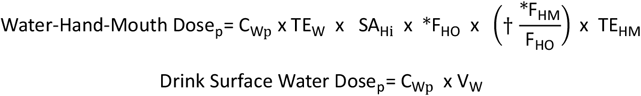

The *p^th^* pathogen is denoted by a subscript of and their concentrations in soil and surface water are denoted as C_s_ and C_Wp_, respectively. * means truncated with a lower bound=0 and upper bound=1, † means the surface area of hand-to-mouth contact cannot exceed the surface area that was contaminated during hand-to-object contact, thus the fraction is truncated at 1.

Fate parameters obtained from the extant literature to estimate exposure to pathogens through indirect contact include: the transfer efficiency of the environmental fomite to the hand (soil: TE_S_, water: TE_W_), the total surface area of the child’s hand (SA_Hi_), the fraction of the child’s hand contacting the environmental object (F_HO_), the fraction of the hand mouthed by the child (F_HM_), and the transfer efficiency of environmental residual from hand-to-mouth (TE_HM_) (Table 1). Total hand surface area (cm^2^) used in this model was based on estimated surface area parameters for children between the ages of six months to less than six years.(31) Standard deviation for total hand surface area (cm^2^) per age category (6 to 11 months, 12 to 23 months, 24 to 35 months, and 36 to 72 months) was calculated by dividing the difference of the EPA-reported mean and 95^th^ percentile for hand surface area by the 95^th^ quantile of a standard normal distribution (1.645).(31) Each age category was equally represented during simulation by sampling the probability of obtaining a random child within each of the unequal month spans and respective hand surface area mean and standard deviation (SA_Hi_). The distribution for the fraction of the child’s hand involved in hand-to-object contact (F_HO_) and hand-to-mouth contact (F_HM_) was calculated by minimizing the squared differences between theoretical and empirical quantiles.(39) Transfer efficiency of environmental residue from hand-to-mouth (TE_HM_) was estimated with a single point estimate due to the lack of literature to infer a distribution for all pathogens used in this analysis.(40) Our exposure model assumes that the hand region that contacted the object is the same region that contacted the mouth. This assumption is supported by the finding that regardless of the type of interaction, hand contact predominantly involves the fingers.(39) When summing across estimated indirect doses, hand size was held constant, while soil adherence and hand area that contacted the object and then mouth varied between summed contact events. To estimate direct ingestion of soil or surface water, the volume of respective substance placed in the mouth during a geophagia (V_S_) or drinking occurrence (V_W_) was estimated with a single parameter because of the lack of literature to describe the distribution of direct ingestion occurrences (Table 1).(14, 26, 32)

## Results

### Correlation in pathogen concentrations

Estimated pathogen concentration distributions (Table 2) were all positively correlated in surface water, but many were not positively correlated in soil (Fig 1). The 95% credible intervals (CI) for the correlations revealed that four pathogen comparisons in surface water were significantly correlated at the site- and neighborhood-level: adenovirus 40/41 vs. EPEC (site CI: 0.05, 0.89; neighborhood CI: 0.17, 0.77), adenovirus 40/41 vs. EAEC (site CI: 0.03, 0.87; neighborhood CI: 0.15, 0.73), ETEC vs. EPEC (site CI: 0.1, 0.86; neighborhood CI: 0.20, 0.74), and EPEC vs. EAEC (site CI: 0.15, 0.88; neighborhood CI: 0.20, 0.75) (S1 Fig). In soil, significant neighborhood-level negative correlation was observed for *Cryptosporidium spp.* vs. adenovirus 40/41 (CI: −0.56, −0.14), and positive correlation for EAEC vs. adenovirus 40/41 (CI: 0.06, 0.61), and EAEC vs. ETEC (CI: 0.3, 0.76). Overall, the strongest pathogen correlations in soil and surface water were observed among human adenovirus 40/41 and pathogenic *E. coli* and among the *E. coli* themselves.

**Table 2.**
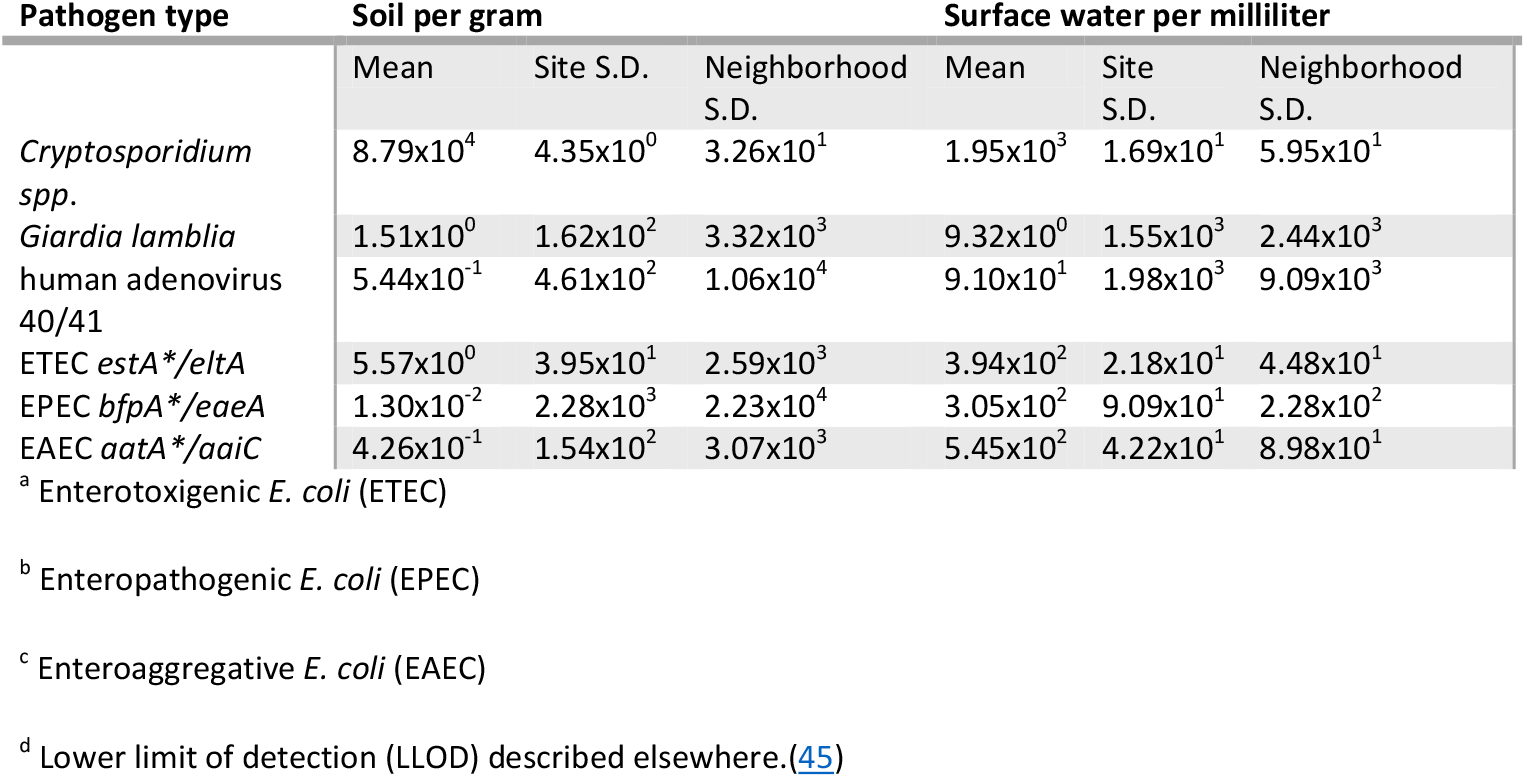
Tobit model-simulated mean concentrations and site-level and neighborhood-level standard deviations (S.D.) of enteric viruses, bacteria, and protozoans detected per gram of soil and milliliter (mL) of surface water public areas of Kisumu where children play.

**Figure 1.**
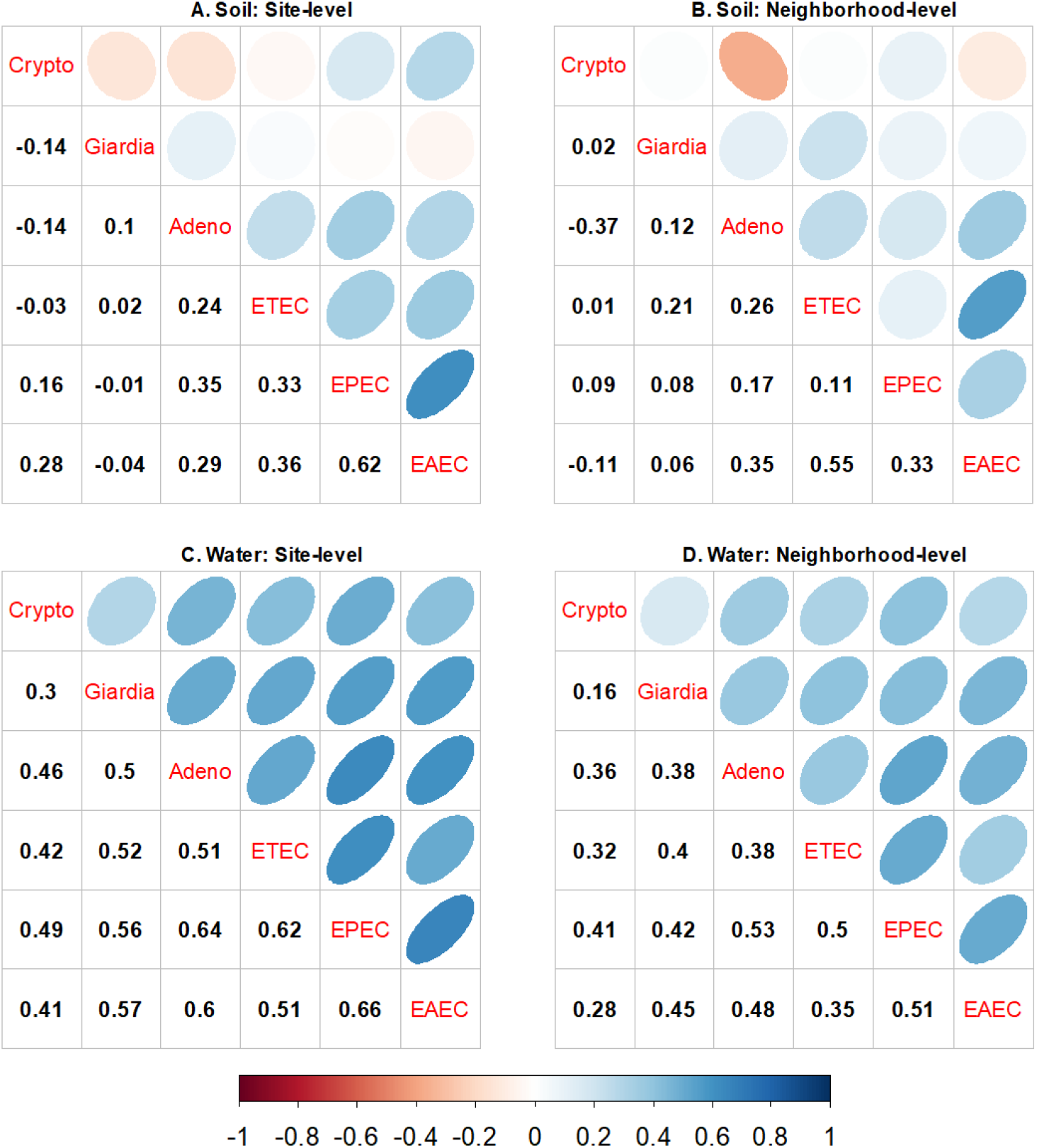
Site- and neighborhood-level correlation in concentration of enteric pathogens in soil and surface water at public areas of Kisumu, Kenya. Negative and positive correlation shown in the lower left quadrants of each grid are reflected in orange-red and blue circles, respectively, in the upper right quadrants of each grid. The highest rho indicated by the darkest color and narrowest shapes.

### Pathogen exposure doses by behavioral pathway and spatial range

Pathogen doses from soil contact (Fig 2) were always lower than doses from surface water contact (Fig 3). All surface water contact resulted in a mean ingestion of at least one of any type of pathogen, with the exception of Giardia for one water-hand-mouth contact. If frequency of contact with soil or surface water is held constant, geophagia or drinking surface water always resulted in higher pathogen doses compared to soil/water-hand-mouth contact (Fig 2C/2D vs. Fig 2A/2B; Fig 3C/3D vs. Fig 3A/3B). However, if hand-mouth contact occurs more often than geophagy or drinking surface water, then doses resulting from hand-mouth contact could exceed exposures from direct ingestion. For example, if a child exhibited ten cumulative soil-hand-mouth contacts and one geophagia contact during play at one site, the EAEC dose for soil-hand-mouth (~80 bacteria, Fig 2A, solid box) would exceed the EAEC dose for geophagia (~1 bacteria, Fig 2C, dashed box). Overall, pathogen doses from contact with soil or surface water increased if children played at multiple sites in the neighborhood, versus just one site (Fig 2B/2D vs. Fig 2A/2C; Fig 3B/3D vs. Fig 3A/3C), but the magnitude of change depended upon the pathogen type.

**Figure 2.**
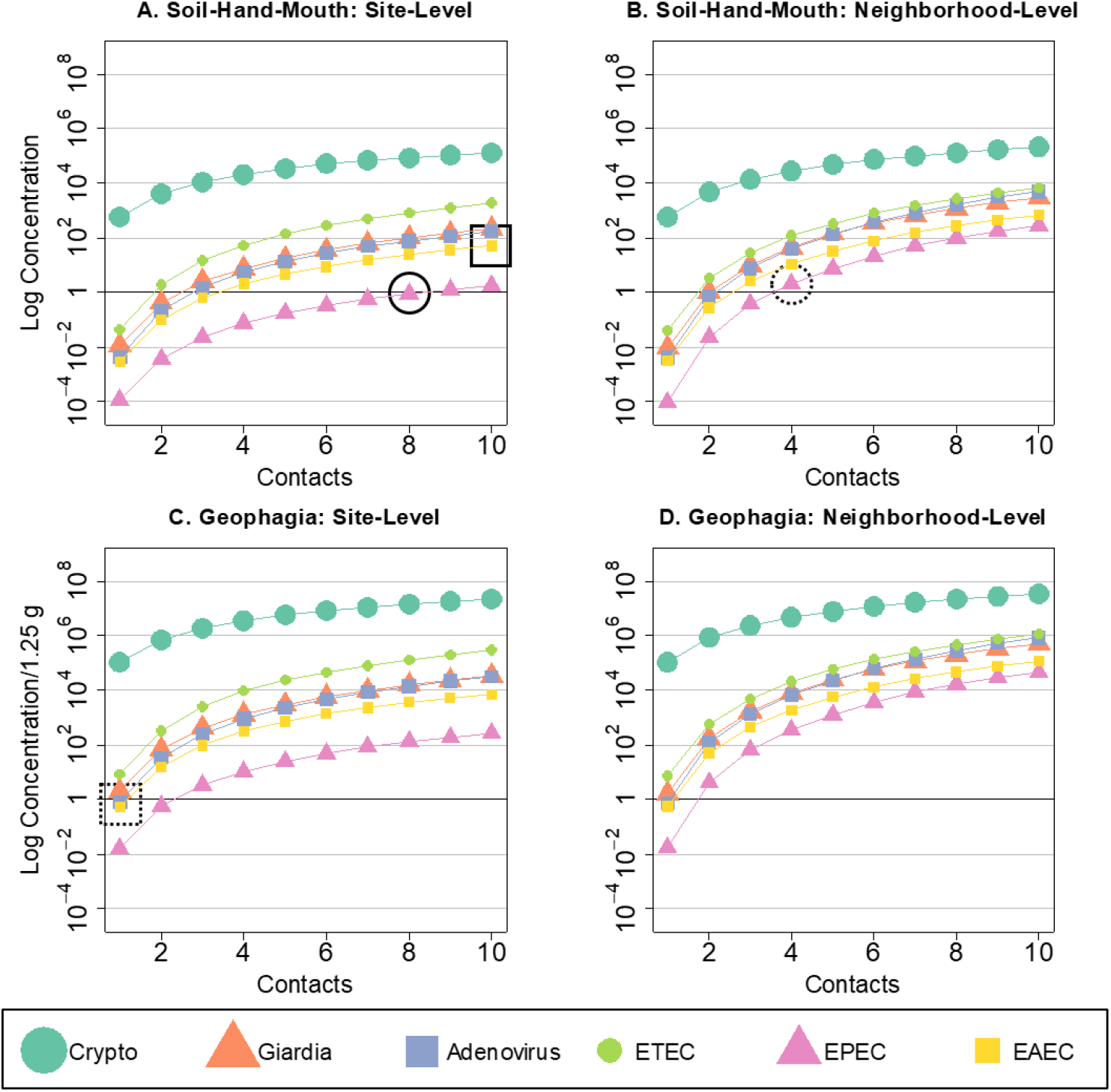
Mean concentration of six enteric pathogens ingested with increased frequency of soil-hand-mouth or geophagy behaviors at site-restricted versus neighborhood levels of spatial scale.

**Figure 3.**
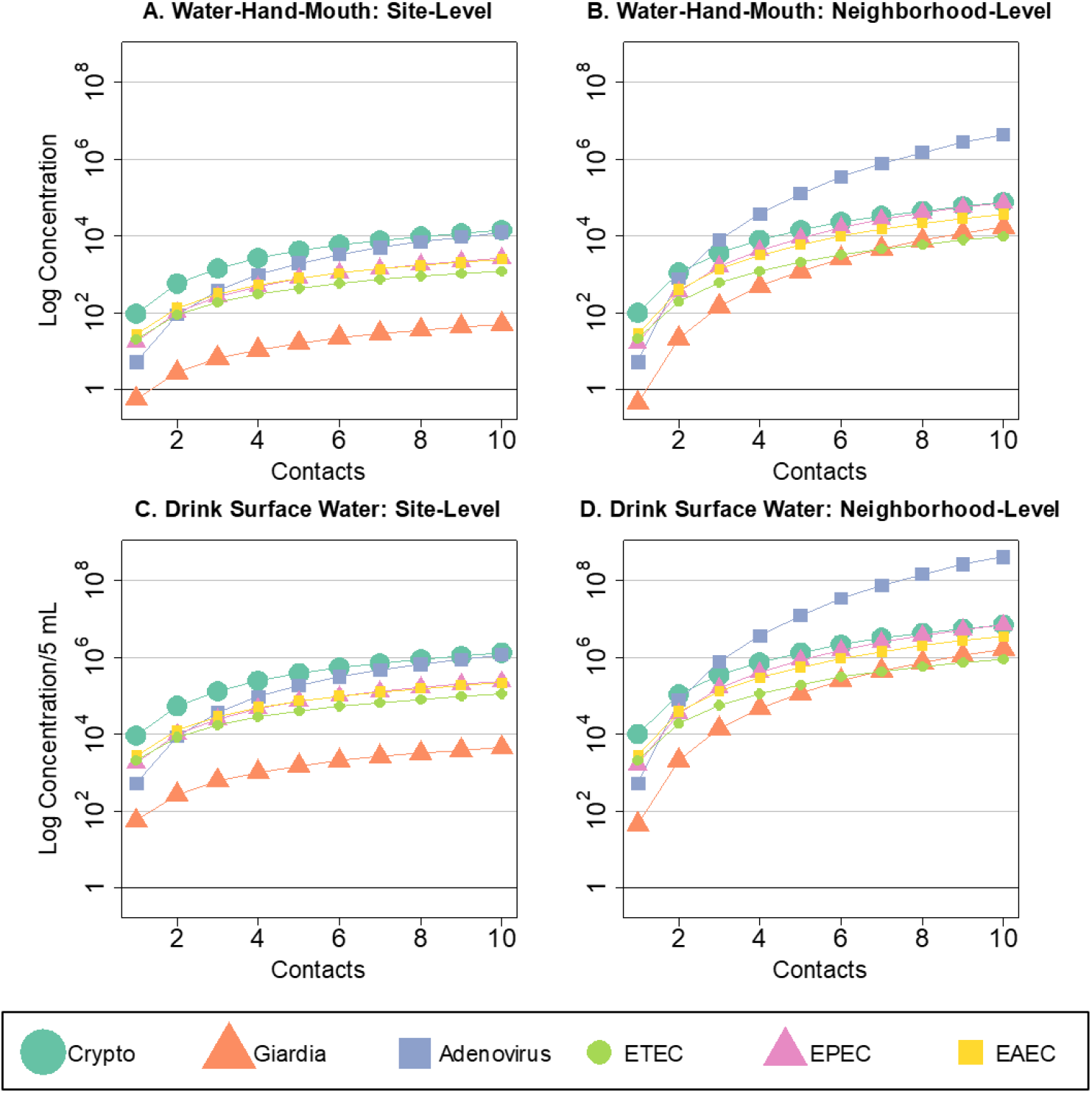
Mean concentration of six enteric pathogens ingested with increased frequency of hand-to-mouth or surface water drinking behaviors at site-restricted versus neighborhood levels of spatial scale.

Pathogen-specific dose distributions from indirect and direct contact with soil and surface water at site- and neighborhood-level are reported in S2-S7 Figs. The mean dose *D*(*k*) of pathogens ingested during contact with soil were largest for *Cryptosporidium spp.* (Fig 2). However, *Cryptosporidium spp.* dose did not considerably increase with increased behavior frequency or increased spatial scale of play from site- to neighborhood-level. The lowest pathogen dose ingested during soil contact was EPEC—yet the dose of EPEC substantially increased for neighborhood vs. site-level play. For example, it required two times more soil-hand-mouth contacts to ingest the same mean dose of EPEC at site-level play (~8 contacts, Fig 2A, solid circle) as neighborhood-level play (~4 contacts, Fig 2B, dashed circle). Most noticeably, the dose of human adenovirus 40/41 from surface water contact exponentially increased as spatial scale expanded from site to neighborhood play and surpassed all other pathogen doses at neighborhood-level exposure.

### Exposure to diverse pathogen types

Fig 4 illustrates the probability of ingesting one or more pathogens for 1 to 10 indirect or direct contact(s) with soil or surface water during site-restricted or neighborhood play, where a successful ingestion is defined as DNA of 1 or more pathogens. Across all behaviors, the probability of ingesting more than one pathogen type intensified as spatial scale expanded from site to neighborhood play. For example, the probability of ingesting six pathogens from two water-hand-mouth contacts during site-restricted play (44%, Fig 4E, solid box) increased by about 16% if the child exhibited the same behavior and frequency during neighborhood-level play (60%, Fig 4F, dashed box). Soil-hand-mouth contact resulted in the lowest risk of ingesting diverse pathogen types compared to all other behaviors practiced at the same frequency. This is especially evident for soil-hand-mouth contact during site-restricted play where the probability of ingesting all 6 pathogens did not exceed 35% for 10 contacts. Any contact with surface water posed a high risk for ingestion of diverse pathogens and is demonstrated by an > 90% probability of ingesting 6 pathogens during ≥5 water-hand-mouth contacts and ≥3 drinking water contacts during neighborhood-level play.

**Figure 4.**
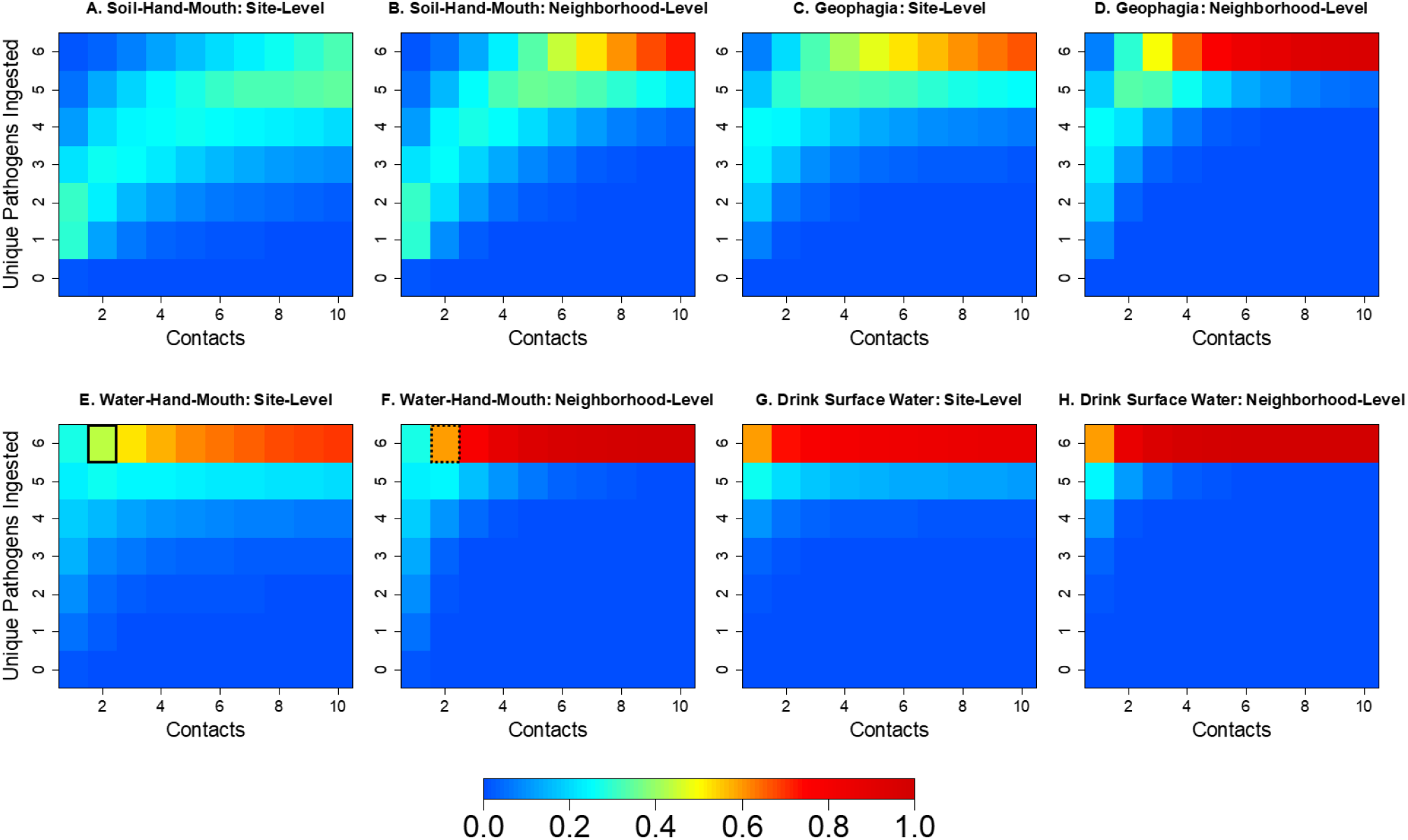
Probability of ingesting one or more enteric pathogens with increased frequency of exposure contact with soil or surface water at site-restricted and neighborhood levels of scale.

## Discussion

Our prior work has demonstrated that < 5 yr. children living in low-income neighborhoods of Kisumu, Kenya are exposed during play in public residential areas to soil and surface water contaminated by human and animal feces and a diverse range of enteric pathogens.(23) This study addresses the next question as to how children are impacted by playing in these settings. Specifically, we quantify the impact of different child exposure behaviors, environmental transmission pathways, and spatial situations on the dose of and diversity in enteric pathogens ingested by young children as a result of contact with public residential areas. When holding frequency of behaviors constant, dose and probability of multiple enteric pathogen exposure were greater when a child ingested: (1) surface water versus soil, (2) greater volumes of soil (geophagy) or surface water (small mouthfuls) versus soil/water-hand-mouth contact, and (3) soil or surface water from multiple neighborhood locations versus just one spatially-restricted site. The evidence that children can be exposed simultaneously to multiple enteric pathogens during some play conditions indicates that some exposure pathways may be more important than others in elevating the risk of infection or even co-infection by different pathogens.

We kept contact frequency of behaviors at a constant rate to examine relative relationships between dose and diversity. In our studies of child play in public areas in Haiti, geophagy was 6 times more common than drinking surface water (0.9/hr vs 0.15/hr), and hand-to-mouth was roughly 9 (child) to 20 (infant) times more common than geophagy.(33) If we assume that child behavior frequencies are generalizable across geographic contexts, then child exposure to low doses (10^0^-10^2^) of at least one type of pathogen from short durations of play outside their home is certain, due to the pervasiveness of pathogen contamination in these neighborhoods and frequent hand-to-soil and hand-to-mouth contacts.(33) Multi-pathogen exposure was common for even the lowest risk behaviors; for example, the summed probabilities for exposure to 2 or more pathogens for one soil-hand-mouth contact during site-level play was 69%, although exposure to all pathogens was unlikely at even 10 contacts (<35%). Yet, contact with surface water was more dangerous, with only a few contacts resulting in exposure to all six pathogens. In reality the rate of different child behaviors vary greatly between children and across populations of children, so the ranking of actual exposure risks associated with different behavioral, environmental, and spatial conditions should be evaluated based upon specific cultural settings. Nonetheless, the threshold of contact between children and public domains in Kisumu for exposure to pathogens is high.

Another notable discovery was that exposure of the child to multiple public locations in their neighborhood, such as with free roaming child play, significantly increased pathogen doses and risk of multi-pathogen ingestion. Consideration of space as a determinant of environmental exposure to fecally-transmitted pathogens is rarely explicitly included as a function of exposure modeling, and when it is, spatial units tend to be defined using households or clusters of households as units of study. Yet, our prior studies suggests that young children in urban, low-income settings – even those less than 24 months - are not immobile elements whose environmental exposures can be defined by these architectural boundaries.(33) While this study provides theoretical insight on potential child exposure conditions if children’s range of movement is considered the unit of study, very little is actually known about how much time children in low income settings spend in public areas and how far they roam.

There were some pathogen-specific differences in exposure dose concentration curves. *Cryptosporidium spp.* was the most common pathogen in soil, at relatively stable concentrations across the neighborhood, which led to higher exposure doses. In this study we did not determine how many of these samples were *C. parvum* or *C. hominus* types, which are typically considered responsible for human infections. Although the public health importance of these exposures remains unclear, the 18S subunit gene indicator used in this study to detect Cryptosporidium was the same indicator used in a study of diarrhea in children less than two years of age in Bangladesh, which found *C. meleagridis* species are also common causes of child infection.(41) Pathogenic *E. coli* dose curves generally behaved similarly to each other, although soil appears to be a less common transmission pathway for EPEC. Human adenovirus and Giardia concentrations were highly varied between sites, compared to other pathogens, which indicates that child exposure to multiple public sites was an important determinant of greater doses.

This study has several important limitations. Concentrations of enteric viruses, bacteria, and protozoan pathogens were estimated by quantitative reverse transcription Polymerase Chain Reaction (qRT-PCR). While use of a multi-pathogen qRT-PCR process reduced methodological sources of variability in concentration estimates, and ergo exposure doses, it does not distinguish between viable and non-viable organisms. Pathogen exposure doses presented in this study may overestimate the number of infectious pathogens children ingest through contact with soil and surface water. There is a lack of information on how well qRT-PCR correlates with other approaches for quantifying pathogen environmental exposure, and with child disease outcomes in settings like Kisumu. However, PCR detection of microbial source tracking (MST) markers in Indian households was associated with diarrhea symptoms in one study.(21) MST markers may occur in the environment more frequently than infectious pathogens, and it is unclear whether MST presence is a reliable proxy for pathogen dose. Validating health outcomes associated with exposure doses in Kisumu children was outside the scope of this study, but is critical to understanding the importance of public exposure pathways in pediatric enteric infections. More information on dose-response in children in low income settings is needed that accounts for potential interdependencies in the presence of pathogens.

The findings of this study provide plausible explanations for why recent large scale trials of household WASH interventions have reported little(42) or no impact on child diarrhea(43) and why the global diarrhea disease burden remains high despite marked improvements in basic water and sanitation access in recent decades.(44) Furthermore, it provides explanations as to why high levels of sanitation coverage may be required to significantly reduce child diarrhea. Child play outside the household is far more common than appreciated(33) and is likely to continue as long as families live in crowded conditions. Children playing unaccompanied or being cared for by older siblings may contribute to hazardous scenarios where children play in neighborhood areas and contact objects that are unsafe or unsanitary.(33) Household WASH interventions do not typically address the topic of where children play, and do not install barriers between children and soil and water contaminated by the feces of one’s neighbors and domestic animals. Thus, while household WASH improvements can reduce child exposure and infection from some pathways, they may not sufficiently reduce exposure across all pathways to observe differences in diarrhea rates.

In conclusion, our results suggest that exposure to public residential areas poses an important risk for enteric pathogen ingestion for children living in low-income settings with poor sanitary conditions, especially when those behaviors are frequent and of high volume, involve contact with surface water, and occur at multi ple locations in the child’s neighborhood. Addressing these exposures will require broader WASH interventions targeting both child play behaviors and the environmental conditions in which play occurs. Future research should address the scarcity of information about spatial patterns of child behavior and the importance of public exposures in child infection outcomes.

## Supplemental information captions

**Table S1.** Summary of pathogens detected in environmental samples in three Kisumu neighborhoods and respective lower limits of detection.

**Appendix S1.** Statistical Analyses and Additional details of the multivariate tobit model including the random effects for spatial dependence and Gibbs sampling algorithm.

**Figure S1.** 95% credible intervals (CI) for site- and neighborhood-level correlation in estimated concentrations of enteric viruses, bacteria, and protozoans in soil and surface water in Kisumu, Kenya.

**Figure S2.** Concentration distribution of Cryptosporidium ingested with increased frequency of soil and surface water contact at site-restricted versus neighborhood levels of spatial scale.

**Figure S3.** Concentration distribution of Giardia ingested with increased frequency of soil and surface water contact at site-restricted versus neighborhood levels of spatial scale.

**Figure S4.** Concentration distribution of human adenovirus 40/41 ingested with increased frequency of soil and surface water contact at site-restricted versus neighborhood levels of spatial scale.

**Figure S5.** Concentration distribution of ETEC ingested with increased frequency of soil and surface water contact at site-restricted versus neighborhood levels of spatial scale.

**Figure S6.** Concentration distribution of EPEC ingested with increased frequency of soil and surface water contact at site-restricted versus neighborhood levels of spatial scale.

**Figure S7.** Concentration distribution of EAEC ingested with increased frequency of soil and surface water contact at site-restricted versus neighborhood levels of spatial scale.

## Acknowledgements

This study was supported by start-up funding from the University of Iowa to PI Baker and a career development award from the University of Iowa Environmental Health Sciences Research Center (NIH P30 ES005605). We appreciate the support of field staff who assisted with the collection of data, development of assays, or data management.

